# *Drosophila* Mutants that Are Motile but Respond Poorly to All Stimuli Tested

**DOI:** 10.1101/045062

**Authors:** Lar L. Vang, Julius Adler

## Abstract

Adult *Drosophila melanogaster* fruit flies were placed into one end of a tube near to repellents (benzaldehyde and heat) and away from the other end containing attractants (light and a favored temperature). They escaped from the repellents and went to the attractants. Five motile mutants that failed to do that were isolated. They did not respond to any external attractants tested or external repellents tested. In addition, they did not respond well to internal sensory stimuli like hunger, thirst, and sleep. The mutants, although motile, failed to respond to stimuli at both 34°C and at room temperature. Some of the mutants have been mapped. It is proposed that the information from the different sensory receptors comes together at an intermediate, called “inbetween” (Inbet), that brings about a behavioral response. The Boss is defined here.

## I. INTRODUCTION

Organisms are constantly exposed to a variety of external attractants and external repellents as well as to a variety of internal sensory stimuli. How organisms respond to these to bring about behavior is a basic question of life.

One approach for discovering how this works is the isolation and study of mutants that fail here. In this report we show that *Drosophila* flies can be mutated in such a way that, although still motile, they no longer respond well to any sensory stimulus tested. This includes various external attractants and various external repellents as well as internal sensory functions like hunger, thirst, and sleep. The isolation of the mutants that failed to respond was carried out at a high temperature (34 degrees C), but they also failed to respond at room temperature. An account of some of this work has appeared (Vang and Adler, 2016). A preliminary report of some of the results has been presented (Adler, 2011; Vang, Medvedev, and Adler, 2012).

It is proposed that information from all the different sensory receptors comes together in the central brain at a newly found intermediate called “Inbetween” (the products of the *inbet* genes), which sends information to bring about a behavioral response. See Figure 1.

**Figure 1.**
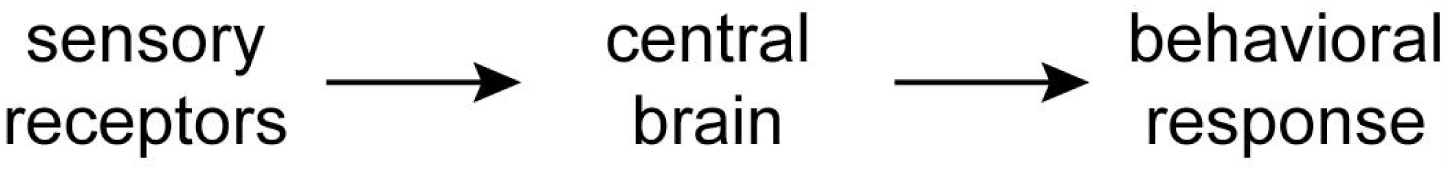
From sensing to response by way of the central brain. Mutations described in this report are located in the central brain.

Work by others has shown that in insects the central brain is a part of behaviors such as courtship (Pavlou and Goodwin, 2013), audition (Clemens et al., 2015), vision (Weir and Dickinson, 2015), and smell and taste (Voshall and Stocker, 2007). The central brain includes the central complex, which is a system of neuropils consisting of the protocerebral bridge, the fan-shaped body, the ellipsoid body, and noduli (Hanesch, Fischbach, and Heisenberg, 1989; Young and Armstrong, 2010; Wolff, Iyer, and Rubin, 2015).

## II. RESULTS

### A. RESPONSES TO EXTERNAL STIMULI

#### 1. RESPONSE TO STIMULI USED TOGETHER

In a 34°C dark room flies were started near two repellents (0.1M benzaldehyde and 37°C) at one end of a tube, away from two attractants (light at 1000 lux and 27°C) at the other end (Figure 2). The parent responded by going away from the repellents and to the attractants (Figure 3A). Mutants that were not motile were rejected, only the motile mutants were studied. This consisted of five mutants, named 1 to 5. Figure 3B shows that such a mutant failed to respond when the four stimuli were together. Each of the five mutants failed to respond to the four stimuli together (Vang and Adler, 2016).

**Figure 2.**
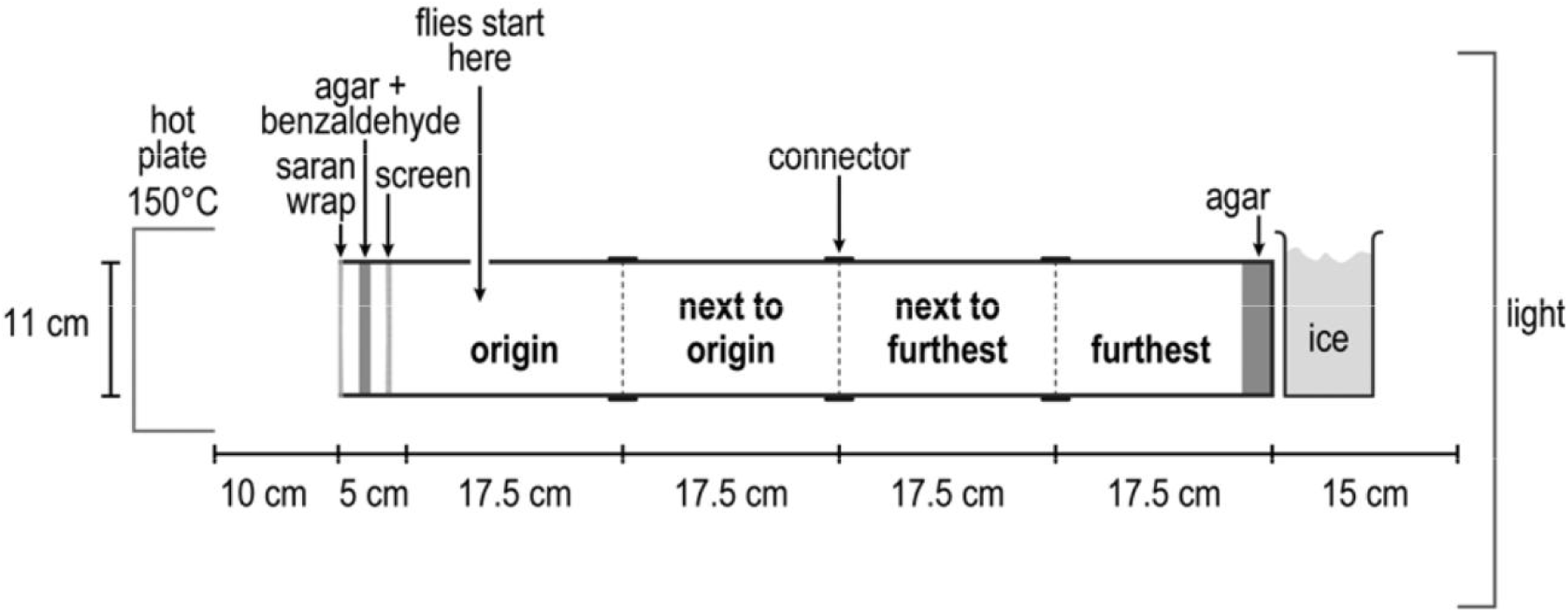
Apparatus for isolating and testing mutants in a 34°C room. At the left end were repulsive 0.1M benzaldehyde and repulsive 37°C (due to a hot plate at 150°C). At the right end were attractive light (1000 lux) and attractive 27°C (due to ice water). The middle was close to 34°C.

**Figure 3.**
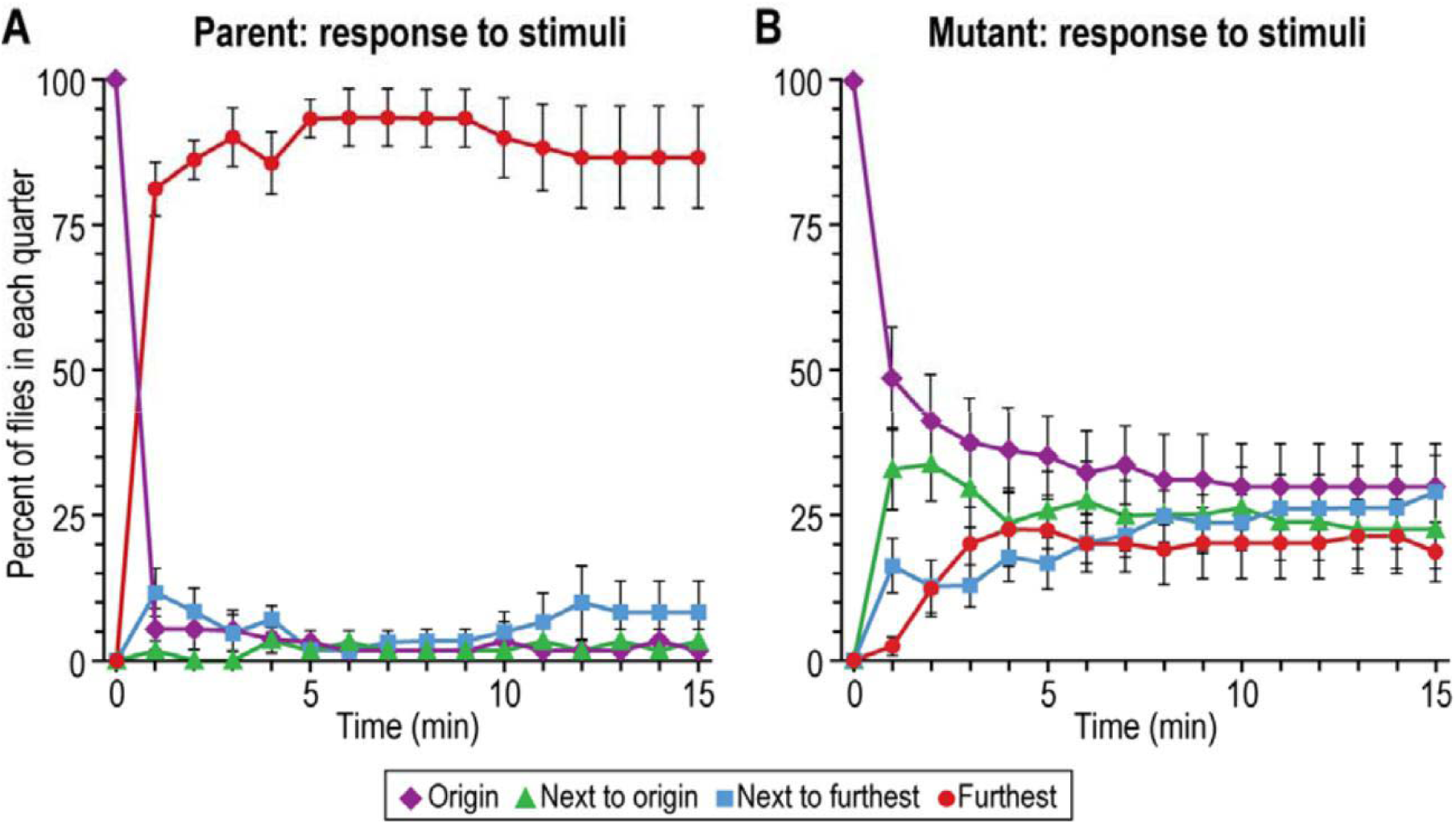
Response to stimuli used together. Repellents (0.1M benzaldehyde and high temperature (37°C) were at the left end, attractants (light, 1000 lux, and a favored temperature (27°C) at the right end. ***A***, Parental response (n=7). ***B***, Mutant 2 (n=8). Flies were tested in a 34°C room with 10 to 20 flies used per trial. Data are mean±SEM.

### 2. RESPONSE TO INDIVIDUAL STIMULI

A single stimulus was presented to flies that were derived from ones that had already experienced the four stimuli used together. For example, the parent went to light only (Figure 4A) while a mutant did not (Figure 4B). Each of the five mutants failed to respond to light only (Vang and Adler, 2016).

**Figure 4.**
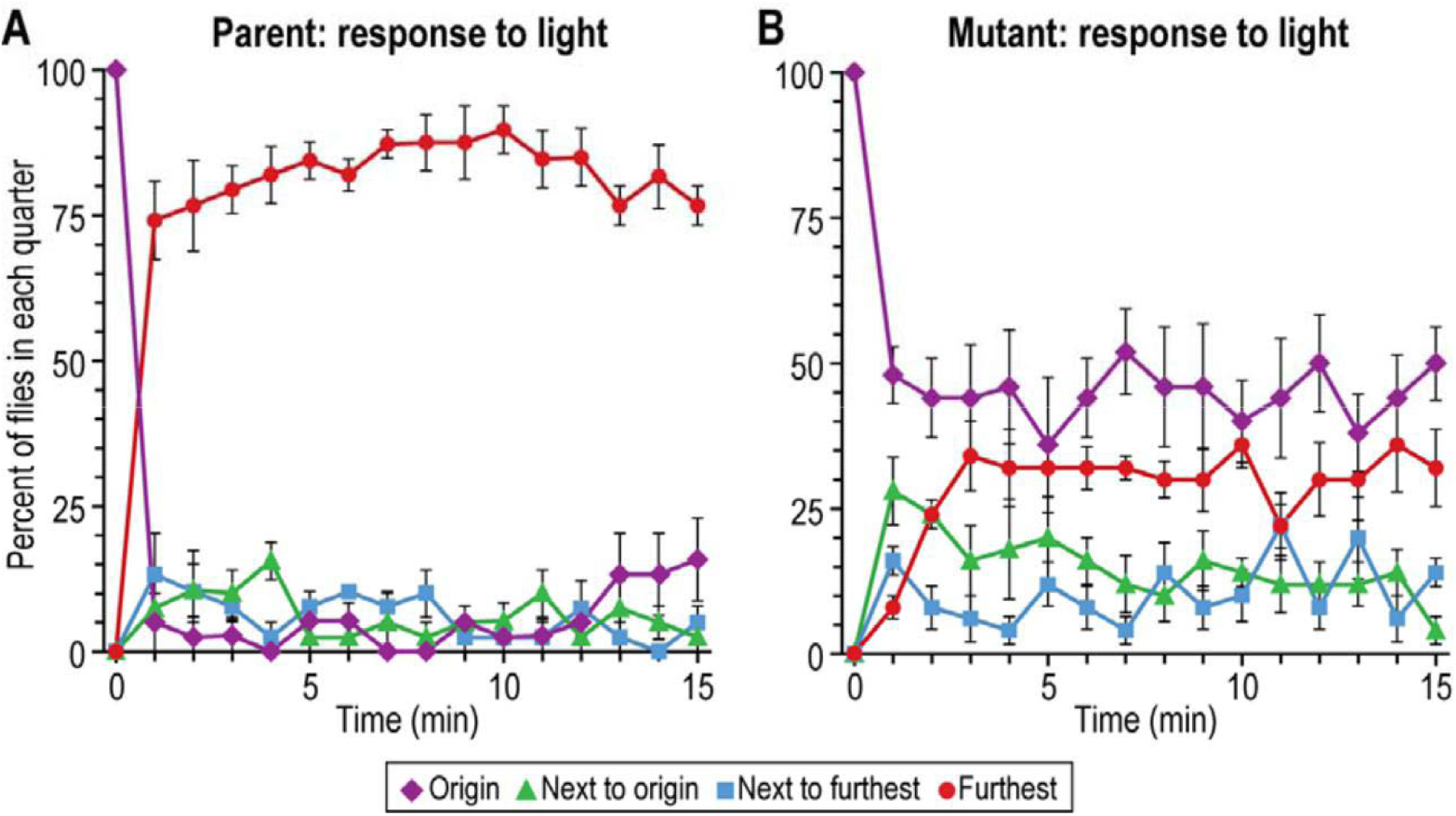
Response to light alone. Light (1000 lux) was placed at the right end as in Fig. 2. ***A***, Parental response (n=4). ***B***, Mutant 1 response (n=5). Flies were tested at 34°C with 10 to 20 flies used per trial. Data are mean±SEM.

For heat alone, the parent was repelled (Figure 5A) but the mutant was not repelled (Figure 5B). That was the case for Mutants 1 and 2 (Vang and Adler, 2016). (The other mutants were not tested for this.)

**Figure 5.**
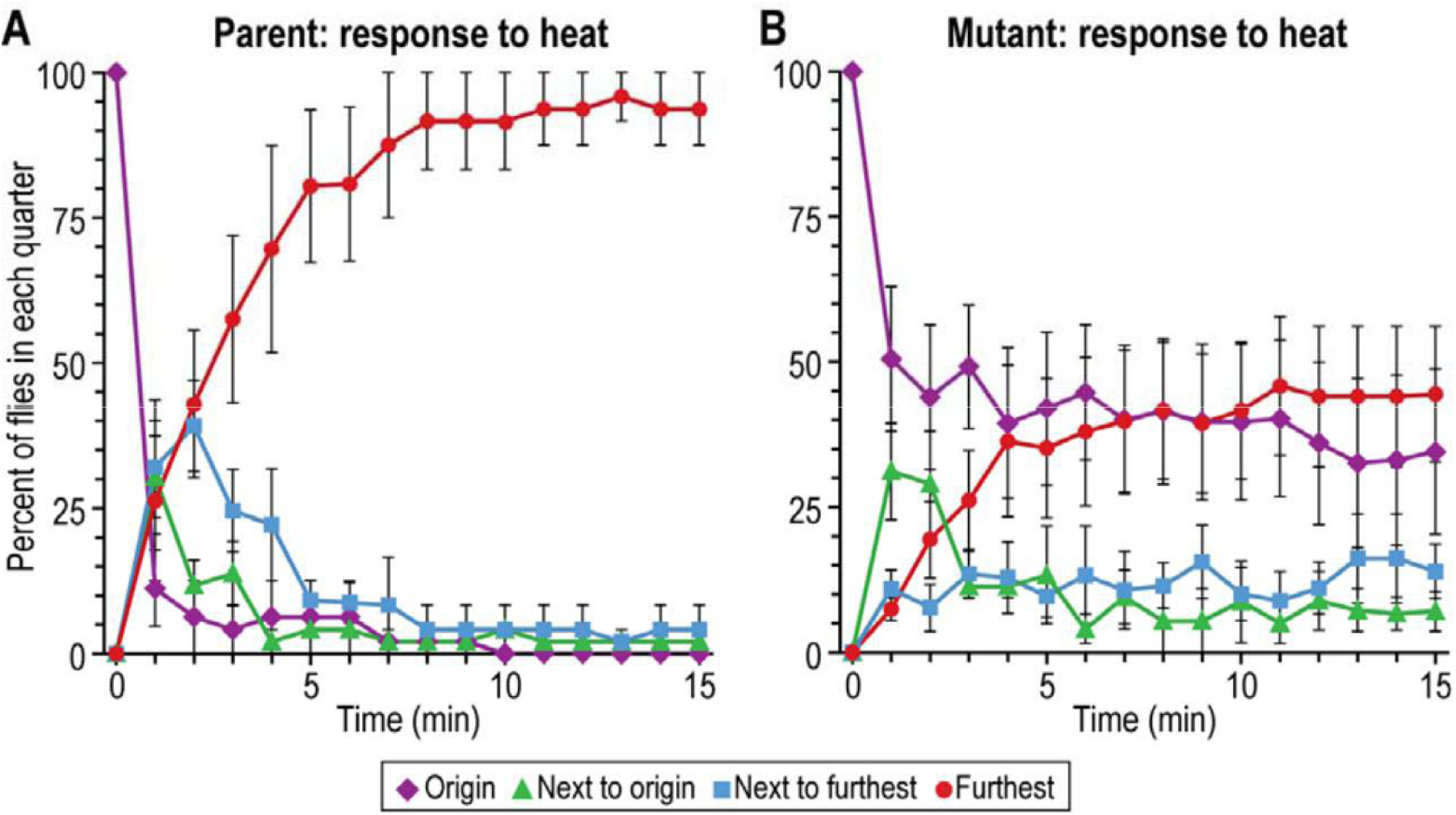
Response to heat gradient alone. The heat source was placed at the left end as in Figure 2. ***A***, Parental response (n=4). ***B***, Mutant 1 response (n=5). Flies were tested at 34°C with 10 to 20 flies per trial. The warm side measured 37°C and the cool side 27°C. Data are mean±SEM.

A similar result was found for benzaldehyde alone: the parent was repelled by benzaldehyde while Mutants 1 and 2 were not repelled. See (Vang and Adler, 2016) for the figures. (The other mutants were not tested for this.)

Thus the mutants were defective not only for the four stimuli used together but also for each stimulus used alone.

#### 3. RESPONSE TO OTHER EXTERNAL STIMULI

These mutants were in addition tested with stimuli that were not among those four used to obtain the mutants:

The mutants were tested for response to the attractant sucrose after starvation (Edgecomb et al., 1994) for 17 to 20 hours. Compared to the wild-type, both Mutants 1 and 2 consumed less sucrose, about 20% as much as the wild-type after subtraction of movement without any added stimuli. See (Vang and Adler, 2016) for the figures. (The other mutants were not tested for this.)

In the case of the repellent quinine, flies were started in a 0.1M quinine half and then they had the opportunity to go into a non-quinine half (see Vang et al., 2012, for details of the method). The parent went into the non-quinine half but Mutant 1 and Mutant 2 did not. See (Vang and Adler, 2016) for the figures. (The other mutants were not tested for this.)

To test response to gravity, these flies were placed into a vertical tube and pounded down, then at every minute the flies in each third of the tube were counted (see Vang et al., 2012, for details of the method). The parent responded by climbing up while Mutants 1 and 2 climbed up 10% as well after subtraction of movement without any added stimuli. See (Vang and Adler, 2016) for the figures. (The other mutants were not tested for this.)

Thus these mutants, isolated by use of the four stimuli, were defective even for stimuli that were not present during their isolation.

#### 4. MOVEMENT WITHOUT ANY ADDED STIMULI

In the absence of any stimulus added by the experimenters, the parent (Figure 6A) and the mutant (Figure 6B) moved similarly, indicating that motility alone is about the same in parent and mutant. This was found also for Mutants 2, 3, 4 and 5 (Vang and Adler, 2016). Aside from our seeing the flies, these results tell that the mutants are motile.

**Figure 6.**
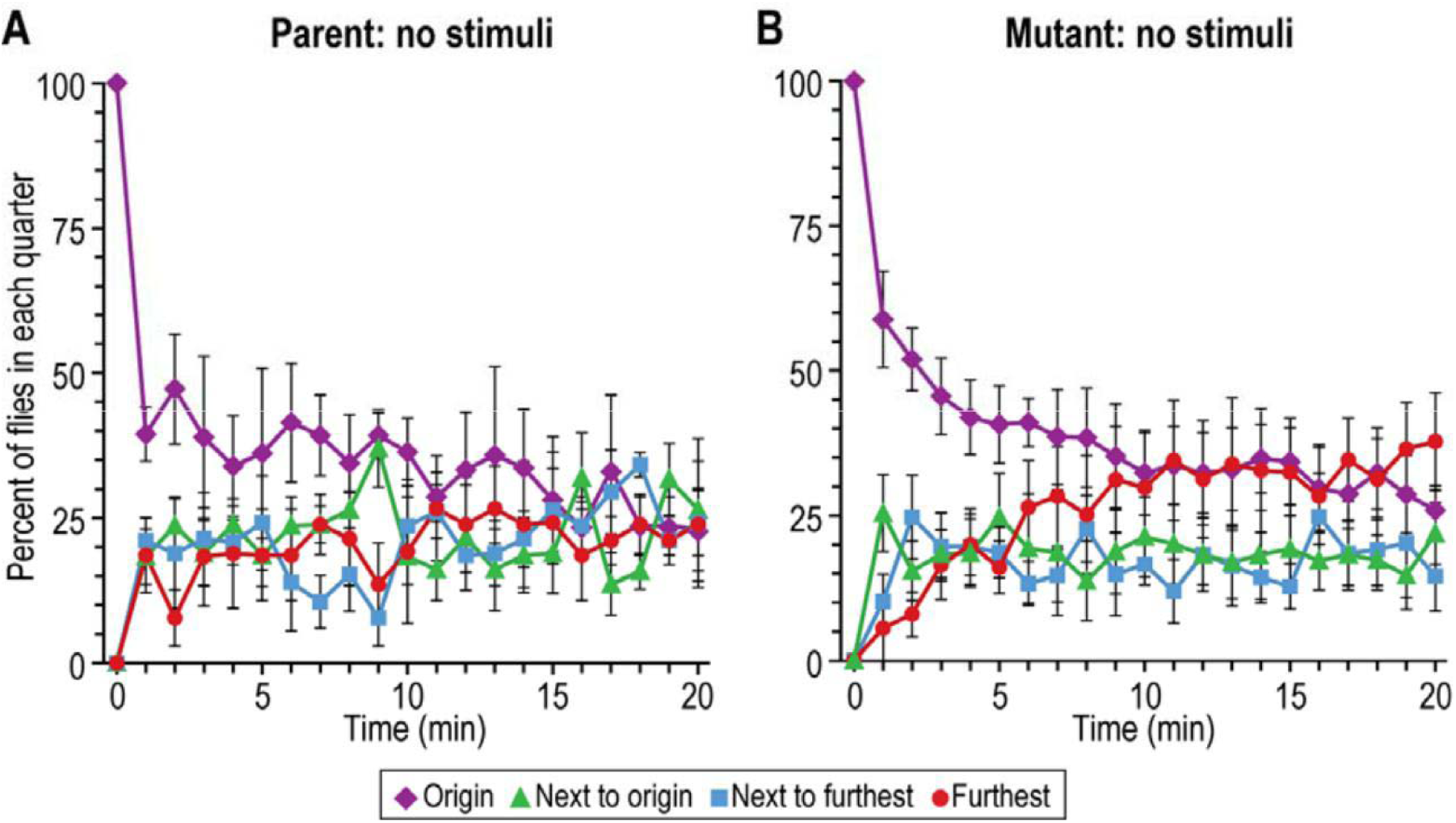
Response without added stimuli**. *A***, Parental response (n=4). ***B***, Mutant 1 response (n=6). Flies were tested at 34°C with 10 to 20 flies used per trial. Data are mean±SEM.

#### 5. EFFECT OF INCUBATION TEMPERATURE

All the work reported above was carried out in a 34°C room in order to allow, if necessary, isolation and study of conditional mutants, i.e. mutants defective at 34°C but not defective at room temperature. We measured response to light (1000 lux) at room temperature (21 to 23°C). The parent responded to light but all five of the mutants failed to respond to light or responded only 10% as well as the parent, just as they did at 34°C (Vang and Adler, 2016). Thus the mutations are not conditional.

Presumably these mutants are defective to all stimuli at room temperature, not just to light. Figures below show defects at room temperature for hunger, thirst, and sleep. Then how could the mutants survive and grow at room temperature? It must be that the mechanism studied here is not an essential one: flies live and reproduce without it.

## B. RESPONSES TO INTERNAL STIMULI

### 1. HUNGER

Here we focus on hunger (Edgecomb et al., 1994; Melche et al., 2007; Fujikawa et al., 2009; Farhadian et al., 2012; Hong et al., 2012; Itskov and Ribeiro, 2012). To measure hunger we used an apparatus (Figure 7), inspired by and modified from an earlier design (Browne et al., 1960), that we described (Vang and Adler, 2016).

**Figure 7.**
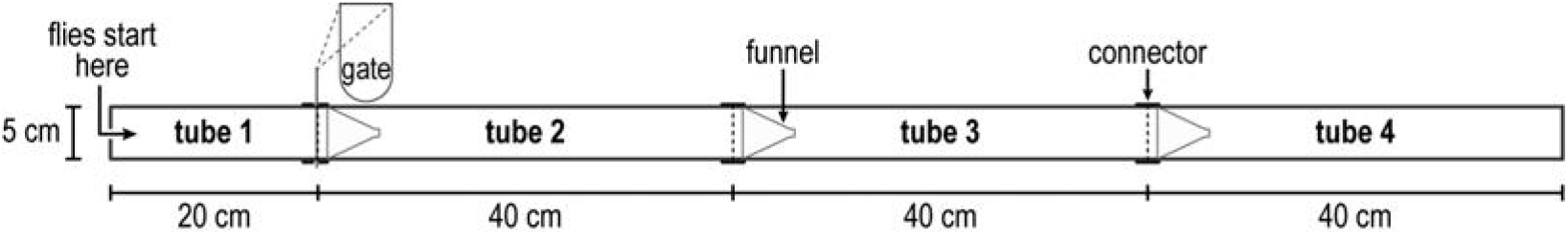
Apparatus for measuring hunger and for measuring thirst. For details see (Vang and Adler, 2016). Tube 1 is called “origin”. Flies were tested at room temperature (21-23°C) for up to 40 hours.

Briefly, in a dark room at 21-23°C male flies – parent or mutants - were transferred into one end (tube 1) of a 5 x 140 cm apparatus containing throughout its length a 5 cm wide strip of wet paper to satisfy thirst but containing no food. Starvation for food began once the flies were put in. Every 10 hours the location of the flies was measured with light on for a few seconds.

At 20 hours the parent had largely left the origin (tube 1) and had begun to accumulate at the end (tube 4) (Figure 8A, solid bars), while the mutant had moved towards the end very little (Figure 8B, solid bars). This is interpreted to mean that the parent is searching for food while the mutant is defective in searching for food.

**Figure 8.**
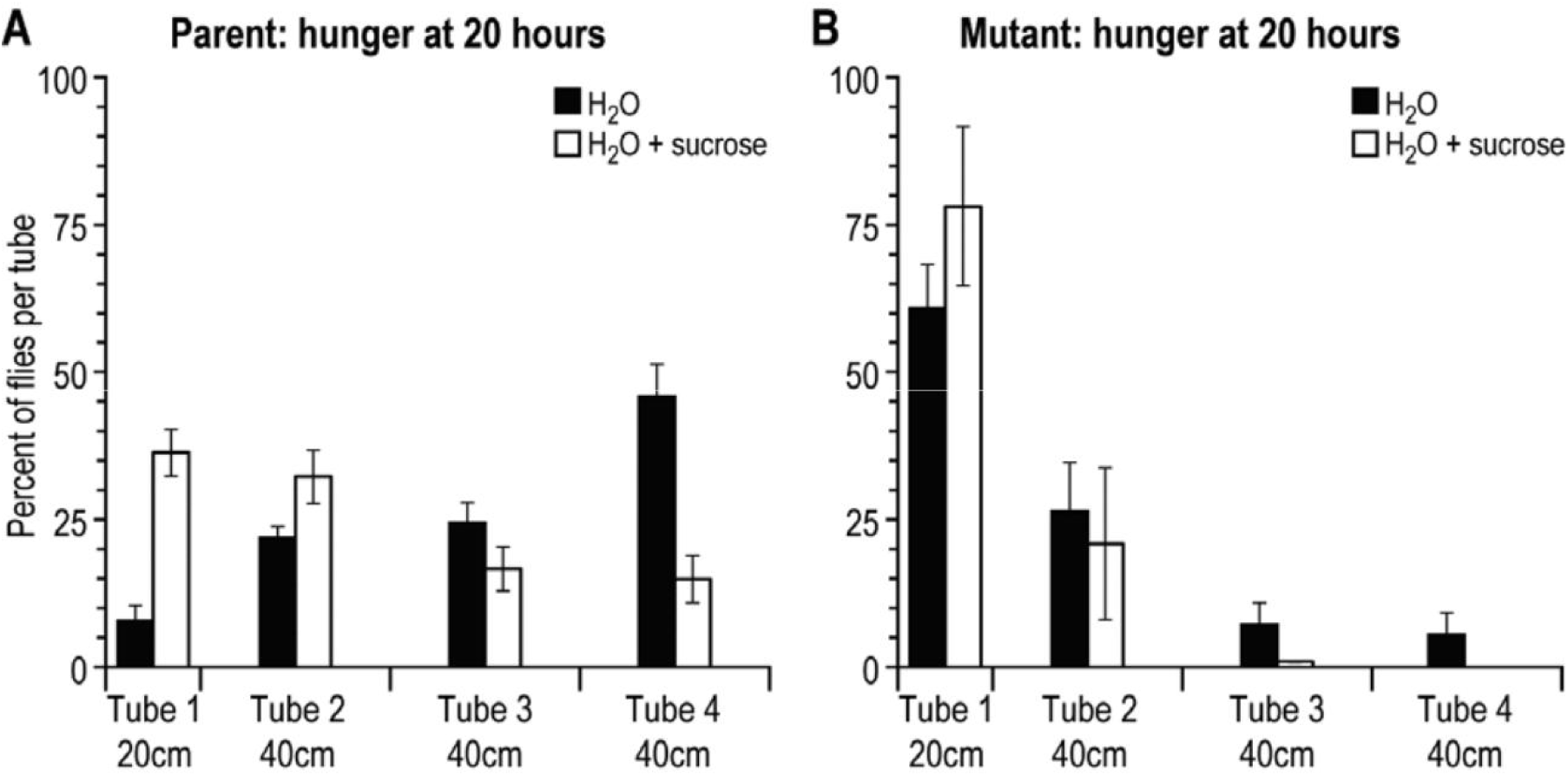
Movement of flies at 20 hours in search for food. Solid: water but no food (no sucrose). Open: water and food (0.1M sucrose). ***A***, Parental response with water only (n=5) and with water + sucrose (n=9). ***B***, Mutant 2 response with water only (n=5) and with water + sucrose (n=4). Data are mean±SEM. See (Vang and Adler, 2016) for Mutant 1; the other mutants were not tested for this. Flies were tested at room temperature (21-23°C) with 40 to 60 flies used per trial.

When food (0.1M sucrose) was added throughout the tube along with the wet strip of paper, the parent moved less far (rather than accumulating at the end) (Figure 8A open bars), while the mutant remained mostly where placed (Figure 8B, open bars). Since sucrose inhibited the movement of the parent, it is supposed that movement without sucrose is due largely to hunger. From these results we conclude that the mutants are defective in hunger.

### 2. THIRST

To study thirst, flies were deprived of water. The procedure is the same as for hunger except that water was omitted and solid sucrose was layered throughout (Vang and Adler, 2016). Mutants 1 and 2 were tested, the other mutants not (Vang and Adler, 2016).

By 30 hours the parent had moved out, presumably to search for water since addition of water inhibited this (Vang and Adler, 2016). The mutant moved out less well than the parent (Vang and Adler, 2016), so we conclude that the mutants are defective in thirst.

### 3. SLEEP-WAKE

The parent and mutants isolated here were studied for sleep and wake according to the procedure of Pfeiffenberger et al. (Pfeiffenberger et al., 2010). The parent was different from the mutants (Figure 9). The parent showed greatest activity at the start and end of the day but not in the middle of the day. Mutant 2 showed high activity throughout the day. Mutant 1 was less active than the parent at the start of the day.

**Figure 9.**
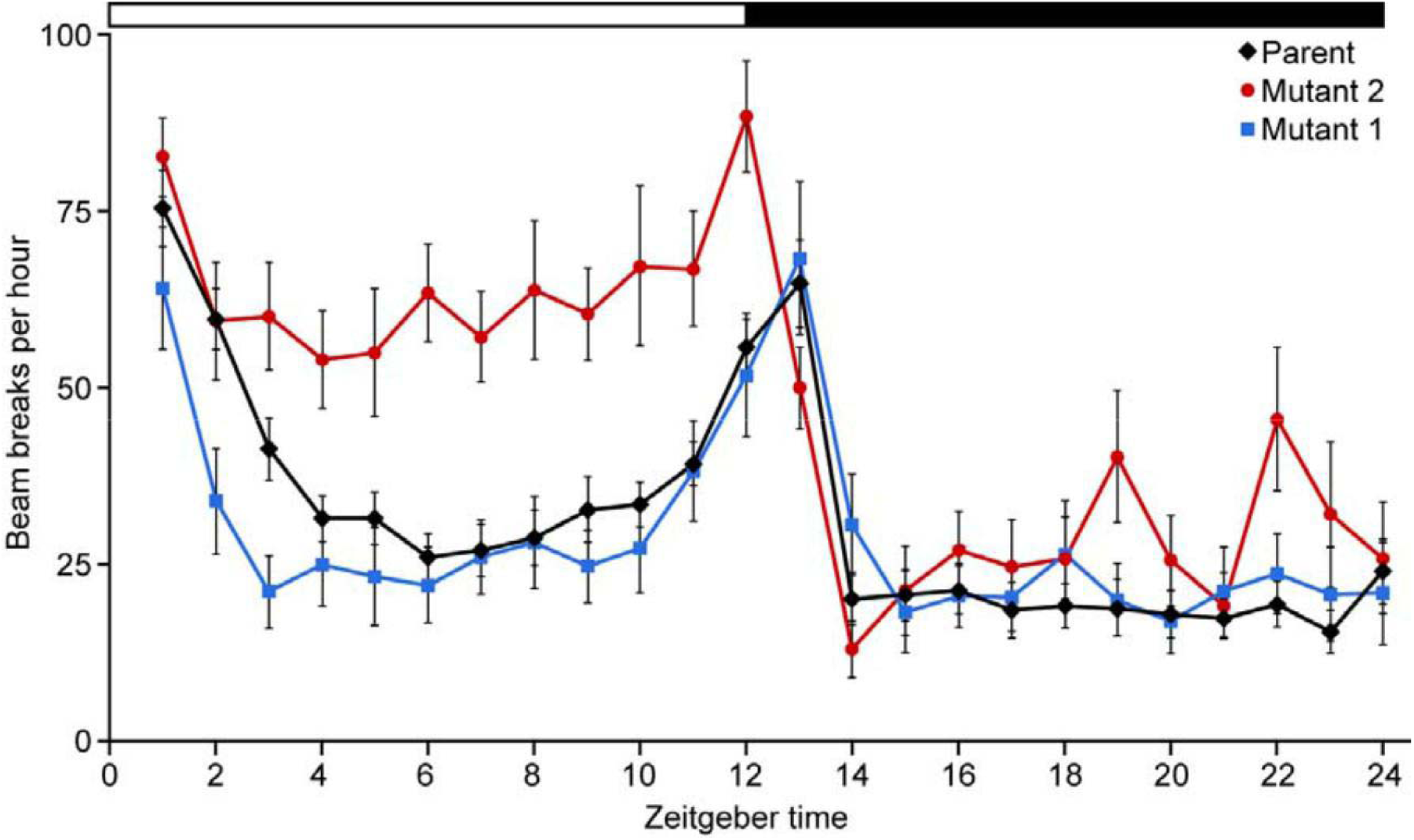
Circadian response. Individual flies are placed into a tube (5 x 20 mm) with an infrared light beam intersecting at the middle of the tube. Mutant 1 (n=24), Mutant 2 (n=24), and parental response (n=24) are recorded over a 24 hour period at 22° C. Data are mean±SEM. There are smaller differences between the parent and the three other mutants (Vang and Adler, 2016).

### C. MAPPING OF THE MUTANTS

We found that Mutant 1 maps in a small gap between 12E3 and 12E5 on the X-chromosome (Vang and Adler, 2016) which we call “*inbetween A”* (*inbetA*). We found that Mutant 2 maps in the *CG1791* gene, a part of the fibrinogen gene of the X-chromosome, or next to it (Vang and Adler, 2016), which we call “*inbetween B*” (*inbetB*).

## III. DISCUSSION

Here we described the isolation and some properties of *Drosophila* mutants that are motile but yet they each fail in response to all external attractants and repellents tested (Figures 3-5) and also they are deficient in response to internal stimuli tested (Figures 8 and 9). Thus, although the mutants are motile, they have:

decreased responsiveness to light
decreased responsiveness to heat and to favorable temperature
decreased responsiveness to repulsive chemicals (like benzaldehyde)
decreased responsiveness to sweet tastants (like sucrose)
decreased responsiveness to bitter tastants (like quinine)
decreased responsiveness to gravity
decreased responsiveness to hunger
decreased responsiveness to thirst
abnormality in some sleep

Because all of these different behaviors are defective, it seems reasonable to say that there is a single place that is responsible, rather than a defect in each of the many different sensory receptors. One possibility for this place is the interaction between sensory receptors and processing of sensory stimuli (Vang and Adler, 2016). Another possibility is that this place is in the central brain (Figure 1) between sensing and response, where all the sensory information comes together at a newly discovered place we call “Inbetween”, the proteins of the *inbetA* and *inbetB* genes.

The mechanism and function of these Inbetween proteins need to be determined further.

An analogy can be made between *Drosophila* mutants missing the part of the central brain studied here and *Escherichia coli* mutants missing their chemotaxis mechanism (Armstrong, Adler, and Dahl, 1967; Parkinson, 1976).

A further analogy can be made with humans. Since the patient’s fall off his bicycle at the age of 10, he had severe epileptic seizures. William Scoville performed experimental surgery to remove the brain’s temporal lobes at the patient’s age of 27. After the operation the seizures reduced drastically but the patient now suffered from amnesia until death at 84. Brenda Milner, a student of Scoville’s, studied the patient: he failed to remember all stimuli (Scoville and Milner, 1957). The patient’s brain after death was examined by histological sectioning, which revealed affected parts in the medial temporal lobes and to a small degree in the orbitofrontal cortex (Annese et al., 2014).

It would appear that all three organisms – bacteria, flies, and people – may have a related mechanism for the path from sensory stimuli to behavioral responses.

In each of these cases, when the behavior was absent the organism was still motile and it still reproduced: thus this behavior is not essential to life, though of course the defective organism is severely handicapped.

Starting in the 1870’s it became apparent to some psychologists that there is a part of the brain, the frontal cortex, that is master of the whole brain; see reviews up to 1970 by the neurophysiologist Aleksandr Luria, who himself modernized this concept and studied syndromes resulting from deficiencies of the frontal cortex (Luria, 1973, 1980). This part of the brain became known as the “central executive” through the research of the psychologist Alan Baddeley (1966) or as the “executive brain” through the research of the neuropsychologist Elkhonon Goldberg (2001), a student of Luria’s. It is now known as “executive control” or “executive function” as well as “prefrontal cortex”. For a review see Goldberg and Bougakov (2007) and Sam Gilbert and Paul Burgess (2008).

The prefrontal cortex of humans and other primates receives information from sensory stimuli recorded in the sensory cortices (visual, auditory, somatosensory, gustatory, olfactory). In addition, the prefrontal cortex receives information from remembered events (memory, emotion) located in the amygdala. Thus these two sources of information merge in the prefrontal cortex to produce a behavioral response. This is described in Helen Barbas’s review (2000), see Figure 10. The prefrontal cortex has also been reported in rodents (Verity Brown and Eric Bowman, 2002; Harry Uylings, Henk Groenwegen, and Bryan Kolb, 2003); and zebrafish (Matthew Parker et al., 2013).

**Figure 10.**
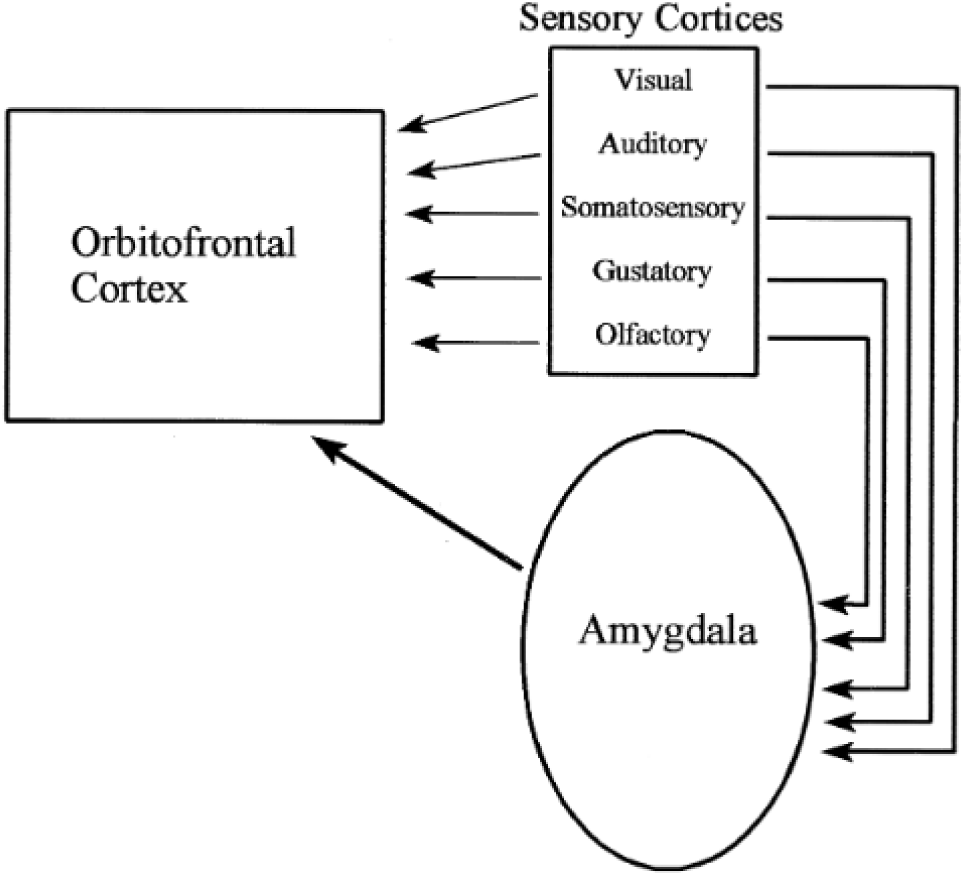
The orbitofrontal cortex (also known as the prefrontal cortex and as a part of the frontal cortex) receives information from sensory cortices and also from the amygdala’s recording of past events and emotions. Modified from Figure 3 of Barbas (2000).

All three parts together – 1. the prefrontal cortex (the orbitofrontal cortex), 2. sensory cortices, and 3. amygdala - constitute what will be called “The Boss”. It is proposed that The Boss in some form occurs throughout biology, in animals, plants, and microorganisms, and that The Boss directs each organism (Adler, 2016). See also Figure 13 and page 60 of Adler (2001) about The Boss. At this time The Boss is an idea without experimental basis.

The control of genes by The Boss could be studied by finding mutants that are in it. Such mutants would be missing the response to all stimuli at a higher temperature but would be normal at room temperature. In the present report Boss mutants did not show up: the mutants described failed at 34°C but they also failed at room temperature. However, only five mutants were studied here; possibly if a much larger number of mutants were isolated there would be some among them that are defective at 34°C but normally responsive at room temperature.

Decision making has been studied in animals, plants, and microorganisms, see Introduction and Results in “Decision making by *Drosophila* flies” (Adler and Vang, 2016, and other references cited). *Drosophila* mutants were isolated by Adler and Vang (2016) that fail in making decisions at 34° C but succeed in making decisions at room temperature, so it seems that The Boss may be in charge of decision making.

The Boss is in control of motivation. “Primary motives are thought to include hunger, thirst, sex, avoidance of pain, and perhaps aggression and fear” according to Charles Cofer and Herbert Petri, 2017. “Motivation is a theoretical construct used to explain behavior. It gives the reason for people’s action, desires, and needs” says Wikipedia (2017). For additional reports about motivation see Vincent Dethier (1976), and Paul Kleinginna Jr and Anne Kleinginna (1981).

Motivation directs behavior, so by being in control of motivation, The Boss becomes in control of behavior. Examples of behavior are eating, drinking, smelling, tasting, seeing, thermosensing, humidity sensing, and gravity sensing, as reviewed by Vang, Alexei Medvedev, and Adler ((2012) and references cited there.

Thus The Boss controls decision making, motivation and behavior.

## IV. METHODS

Details of methods used here are found in in the previous paper (Vang and Adler, 2016): A. Isolation of mutants. B. How to study response to external stimuli. C. How to study response to internal stimuli.

## ACKNOWLEDGEMENTS

Julius Adler is grateful to the Camille and Henry Dreyfus Foundation for six years of grants in support of 31 undergraduate research students named in Adler and Vang, 2016, and in Vang and Adler, 2016. Lar Vang is currently associate research specialist in the Adler laboratory. Robert A. Kreber, a research specialist in Barry Ganetzky’s laboratory, has helped us in studies of the genetics of our mutants. We are grateful to Erin Gonzales and Jerry Yin for showing us how to use the *Drosophila* activity monitory system. Julius Adler thanks Barry Ganetzky for teaching him about fruit flies. Thanks to Millard Susman for criticism. We are thankful to Laura Vanderploeg for the art work.

